# Quantifying and Predicting the Difficulty of Multiple Sequence Alignment with AlDiScore

**DOI:** 10.64898/2026.05.29.727837

**Authors:** Mattis Bodynek, Lucía Martín-Fernández, Julia Haag, Ben Bettisworth, Alexandros Stamatakis

## Abstract

Multiple Sequence Alignment (MSA) constitutes an important and frequent operation in molecular sequence data analysis. There exist numerous tools, algorithms, and criteria to infer an MSA. This plethora of available approaches to MSA may induced an ensemble of divergent MSAs for the same underlying unaligned sequence set. Even a single MSA tool may infer distinct MSAs when varying the input parameters. Hence, when using a diversified set of MSA algorithms and parameterizations, the observed dispersion within an MSA ensemble expresses the difficulty of inferring a robust alignment. We refer to this notion as MSA difficulty. As downstream analyses heavily rely on the MSA, characterizing MSA difficulty for a given unaligned sequence set is critical.

Initially, we show that measures of dispersion within diversified MSA ensembles can reliably predict MSA difficulty. We then assess the adequacy of these measures by computing the average reference-based distance between the MSAs in the MSA ensemble and its corresponding structural reference MSA and subsequently comparing this distance to the corresponding reference-free average distance over all MSA pairs in the ensemble. We find that Blackburne and Whelan’s *d*_pos_ alignment metric is most appropriate as its reference-free 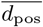 counterpart most accurately approximates the reference-based difficulty computed on BAliBASE reference data. We therefore use 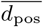 to quantify MSA difficulty on a scale from 0 (easy) to 1 (difficult).

Next, we introduce the `AlDiScore` open-source tool, which uses machine learning to directly and reliably predict reference-free difficulty scores from unaligned sequence sets to completely omit expensive MSA computations. The underlying regression model relies upon a large set of features, including sampling-based measures of transitive consistency. We trained our `AlDiScore` model on a diverse collection of empirical datasets from BAliBASE, TreeBASE, and published studies. Subsequently, we demonstrate that `AlDiScore` attains an *R*^2^ of 0.89 and of 0.84 on unseen AA and DNA sequence sets extracted from the PANDIT v17 database. Finally, we show that there is no correlation between MSA difficulty and the corresponding phylogenetic difficulty of the respective MSA.

## 1 Introduction

Multiple Sequence Alignment (MSA) constitutes a fundamental step in molecular sequence analysis. MSA is generally challenging as the operational unit of an MSA, the homologous site, is difficult to properly define (Roffé 2020). In principle, a site in the genomes of typically distinct species is homologous if it has undergone the “same” evolutionary history over time, and can therefore be matched (aligned) across genomes. However, the inferred MSA substantially depends on the exact definition of homology being used. Even once a definition of homology has been established, *observing* or *inferring* these homologies remains challenging. For instance, homologous sites are often inferred via algorithms that partially disregard the exact evolutionary process that gave rise to them, typically for the sake of computational efficiency and because the underlying true evolutionary history is not known and requires an MSA to at least be estimated in the first place (however, see Redelings and Suchard (2005) and Löytynoja and Goldman (2008) for phylogeny-based approaches to MSA). If we can only estimate, but not observe, homologous sites, verifying the corresponding estimates becomes challenging. One may rely on manually curated structural MSAs, such as those in BAliBASE (Bahr et al. 2001; Thompson et al. 2005). However, such MSAs are rare and labor-intensive to generate.

Nonetheless, inferring homologous sites to generate MSAs is critical to molecular sequence analysis. For example, in phylogenetic inference, the MSA is generally the *only* input data provided to a phylogenetic inference tool (Kozlov et al. 2019; Minh et al. 2020). Due to this dependence on the given input MSA, phylogenetic inference tools may infer different trees on distinct input MSAs for the same underlying sequence set (Wong et al. 2008).

Besides the aforementioned epistemic issues, there also exist practical considerations. Most importantly, inferring an MSA on an unaligned sequence set is computationally expensive. Therefore, widely used MSA tools (Katoh et al. 2002; Sievers et al. 2011; Edgar 2022) typically implement heuristics that exhibit a “good” trade-off between accuracy and run-time. Evidently, these trade-offs may introduce errors into the MSA. In conjunction with distinct underlying alignment models as well as homology assumptions, these heuristics and the corresponding tools implementing them may infer different MSAs on the same input sequence set (Landan and Graur 2007; Wu et al. 2012), that can potentially yield contradictory evolutionary histories in the downstream phylogenetic analyses (Wong et al. 2008). To mitigate this challenge the Muscle v5 (Edgar 2022) alignment tool now offers strategies to infer an MSA ensemble from a unique unaligned sequence set using different alignment parameters. Alternatively, the results of multiple independent alignment methods can be merged into a single consensus MSA using M-Coffee (Wallace et al. 2006), for instance. Efforts to assess alignment accuracy have relied upon generating an MSA ensemble and subsequently rank the distinct alignments in the ensemble (Wong et al. 2008; Shrestha and Adhikari 2022; Serok et al. 2026). Using accurate MSAs improves the accuracy of phylogenetic reconstructions (Serok et al. 2026) and predicted structures (Shrestha and Adhikari 2022).

However, generating an MSA ensemble and subsequently also conducting downstream analyses (e.g., phylogenetic inferences) on all individual MSAs in that ensemble will even be more computationally expensive than the inference and downstream analysis of a single MSA. This is because inferring an MSA ensemble duplicates much of the work required to construct a single MSA. Therefore, the computational cost tends to scale linearly with the ensemble size. Given the prolegomena, it is therefore essential to know, *a priori*, if generating and propagating such an ensemble to the downstream analyses is indeed required for a given unaligned sequence set.

### 1.1 Defining Alignment Difficulty

To decide when such computationally expensive approaches should be employed, we need to quantify the dispersion of an MSA ensemble for a given unaligned sequence set. We refer to this as the “MSA difficulty” of an unaligned sequence set. This concept is closely related to the notion of MSA *reliability*, defined by Herman et al. (2015) which reflects the degree to which a particular MSA can be trusted as an accurate representation of homology among the sequences. This difficulty is inherent to the complexity and dissimilarity of the unaligned sequences (Rost 1999), which can induce multiple plausible, yet conflicting MSAs.

The concept of alignment difficulty was first introduced by Lassmann (2005). They quantified difficulty by how dispersed alignments are in the space of all plausible solutions. In simple alignment cases, distinct alignment programs produce similar alignments, while in difficult cases, the alignments tend to vary to a substantially greater extent. Consequently, MSA difficulty can be empirically observed by considering the level of disagreement among alternative MSAs in an ensemble of MSAs. Throughout the remainder of this paper, we use the term MSA ensemble to denote a collection of alternative MSAs generated via different MSA methods and distinct method parameterizations on the same underlying unaligned sequence set. Greater disagreement within an MSA ensemble indicates higher alignment uncertainty and, therefore, increased MSA difficulty.

For a given MSA ensemble, we can quantify the alignment difficulty by computing a similarity measure between each alternative MSA and an external reference MSA, if available. Thereby, we capture the dispersion of the ensemble around the reference. Such reference-based dispersion scores are highly predictive of MSA error with respect to a reference MSA and have long been used to assess MSA quality (Thompson et al. 1999; Penn et al. 2010). Benchmark datasets, such as BALiBASE (Thompson et al. 2005), provide high-quality reference alignments that are well suited for this purpose (Iantorno et al. 2014).

However, existing benchmark datasets are typically context-dependent and fail to adequately capture the full diversity and complexity of real-world MSA challenges (Iantorno et al. 2014). To quantify the MSA difficulty of a set of sequences *without* a corresponding reference MSA, we first investigate and compare different ensemble-based scores in Section 2.3. Then, we provide a rationale for choosing the 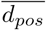 2 metric as the most adequate one to quantify MSA difficulty in Section 2.3.1.

Generating MSA ensembles even for moderately sized datasets is computationally expensive. Consequently, quantifying MSA difficulty for a set of unaligned sequences by inferring such an ensemble is not practical and may also not be necessary, particularly so, for sequences that are “easy” to align.

To address this challenge we introduce the `AlDiScore` open-source tool. `AlDiScore` predicts the MSA difficulty for an unaligned sequence set via a machine learning model that we trained on a large set of MSA ensembles and corresponding difficulties on empirical sequence sets.

In Section 2.4.1 we describe how we generate difficulty labels using 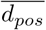 2 on our training data. Thereafter, we describe our feature design process (Section 2.4.2) and how we train the final `AlDiScore` model (Section 2.4.3). Subsequently, we show that `AlDiScore` can accurately predict the difficulty of a putative alignment on *entirely unseen* Pandit data with an *R*^2^ of 0.89 on unaligned AA sequence sets and of 0.84 on unaligned DNA sequence sets (Section 3.2. We also show that `AlDiScore` is up to almost three orders of magnitude faster in predicting the difficulty on large sequence sets compared to directly computing the difficulty by first inferring a corresponding MSA ensemble (Section 3.3). Finally, we discuss additional insights regarding differences in prediction feature importance between AA and DNA data in Section 3.4 and the relationship, or rather absence thereof, between MSA difficulty and the corresponding phylogenetic difficulty (Section 3.5).

## 2 Methods

We initially describe the datasets we used to quantify and predict alignment difficulty in the subsequent subsection. Thereafter, we outline how we compute difficulty labels, how we engineer our features, and how we train `AlDiScore` to predict the MSA difficulty on unaligned sequence sets.

### 2.1 Data

In the first method development step, we use an appropriate BAliBASE subset that comprises structural reference MSAs to determine the most appropriate reference-based as well as reference-free metric for quantifying MSA difficulty on MSA ensembles. In the second method development step we use a collection of 9651 unaligned sequence sets from diverse sources to train our `AlDiScore` predictor. Finally, in the results section, we use a set of entirely unseen sequence sets to assess the accuracy of our `AlDiScore` prediction.

#### 2.1.1 Selected BAliBASE Datasets for Evaluating Difficulty Metrics

We use BAliBASE version 3 (Thompson et al. 2005) as it comprises structural reference “ground truth” MSAs. More specifically, we only use the RV11 and RV12 reference sets (Table 1). The RV11 set contains 74 sequence sets with highly divergent sequences (that is, sequence identity below 20%) that are generally hard-to-align. The RV12 reference set contains 88 amino acid sequence sets with sequence identities between 20% – 40%, which are easier to align. For the sake of simplicity, we refer to RV11 as hard and to RV12 as easy. This selection of subsets allow us to assess the adequacy of candidate metrics for quantifying MSA difficulty on intuitively easy- and intuitively difficult-to-align sequence sets. RV11 and RV12 both comprise full-length sequence sets as well as *truncated* sequence sets that only contain the homologous regions. We expect the full-length sequence sets to harder to align than the curated homologous regions.

**Table 1:** Descriptive statistics for the BAliBASE (Thompson et al. 2005) RV11 (hard) and RV12 (easy) sequence sets. Column *#Sets* contains the number of unaligned sequence sets per partition type. We distinguished between full length and *truncated* sets. Column # *Seqs* indicates the mean number of sequences per sequence set. Column *Length*provides the mean length of the longest sequence within each set. Column 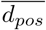 indicates the mean value of the reference-free difficulty label 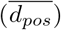 computed on corresponding MSA ensembles (see Section 2.4.1) for details).

| Source | Partition | Identity | # Sets | Type | # Seqs | Length | $\overline{d_{pos}}$ | Reference |
| --- | --- | --- | --- | --- | --- | --- | --- | --- |
| BALiBASEv3 | RV11 | < 20% | 38 | Full Length | 6.87 | 412.29 | 0.51 | (Bahr et al. 2001) |
|  |  |  | 38 | Truncated | 6.87 | 255.71 | 0.40 |  |
|  | RV12 | 20–40% | 44 | Full Length | 9.00 | 585.09 | 0.21 |  |
|  |  |  | 44 | Truncated | 9.00 | 327.75 | 0.14 |  |

**Table 2:** Description of the training sequence sets and their sources. In total, we use 9651 sequence sets, comprising 3851 AA sequence sets and 5800 DNA sequence sets. Column *#Sets* states the number of unaligned sequence sets per data source. Column *# Seqs* indicates the mean number of sequences per sequence set. Column *Length* shows the mean length of the longest sequence within each set. Column 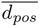 indicates the mean value of the reference-free difficulty label 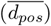 computed on corresponding MSA ensembles (see Section 2.4.1) for details).

| Source | # Sets | Type | # Seqs | Length | $\overline{d_{pos}}$ | Description | Reference |
| --- | --- | --- | --- | --- | --- | --- | --- |
| HOMSTRAD | 193 | AA | 5.45 | 214.41 | 0.07 | Data from HOMSTRAD, used for benchmarking in Formatt. | (Stebbins and Mizuguchi 2004; Daniels et al. 2012) |
| SABmark-Twilight | 158 | AA | 8.33 | 217.08 | 0.31 | Data from the Twilight subset of SABmark, used for benchmarking in Formatt. | (Van Walle et al. 2005; Daniels et al. 2012) |
| SABRE | 272 | AA | 5.72 | 208.33 | 0.42 | Compiled by Robert Edgar. Consistent multiple alignments constructed from SABMARK v1.65. | (Van Walle et al. 2005; Edgar n.d.) |
| BALIiBASEv3 | 386 | AA | 28.71 | 575.18 | 0.27 | Compiled by Robert Edgar. BALiBASE v3. | (Thompson et al. 2005; Edgar n.d.) |
| PREFABv4 | 1665 | AA | 45.32 | 289.61 | 0.18 | Compiled by Robert Edgar. PREFAB v4. | (Edgar 2004; Edgar n.d.) |
| OXBench | 274 | AA | 8.33 | 146.74 | 0.10 | Compiled by Robert Edgar. OXBench. | (Raghava et al. 2003; Edgar n.d.) |
| Arthropods-P450s | 281 | AA | 35.48 | 530.17 | 0.04 | Data from the cytochrome P450 gene family (CYP genes) of arthropods | (Charamis et al. 2025) |
| TreeBASEv1 - AA | 622 | AA | 48.51 | 767.20 | 0.10 | Sampled from TreeBASE | (Piel et al. 2000) |
| TreeBASEv1 - DNA | 2497 | DNA | 73.42 | 2265.78 | 0.06 | Sampled from TreeBASE | (Piel et al. 2000) |
| BALIiBASE (mdsa_all) | 498 | DNA | 34.84 | 1857.66 | 0.50 | Back-translated AA datasets using all tBLASTn hits with an E-value score above the threshold of 0.001. | (Carroll et al. 2007) |
| OXBench (mdsa_all) | 276 | DNA | 10.64 | 441.83 | 0.19 | Back-translated AA datasets using all tBLASTn hits with an E-value score above the threshold of 0.001. | (Carroll et al. 2007) |
| SMART (mdsa_all) | 690 | DNA | 63.77 | 455.74 | 0.31 | Back-translated AA datasets using all tBLASTn hits with an E-value score above the threshold of 0.001. | (Carroll et al. 2007) |
| RobLanf | 1839 | DNA | 109.48 | 360.08 | 0.07 | Single-locus DNA alignments | github:roblanf/BenchmarkAlignments |

#### 2.1.2 Training Data

We assembled a diverse collection of 9651 unaligned sequence sets to train and initially validate our model. Of these, 3851 are amino acid (AA) datasets and 5800 are DNA datasets. We obtained these sequences by unaligning publicly available MSAs. Corresponding summary statistics and publication sources are listed in Table 2.1.2. Note that the majority of the servers where the databases were originally published have been deprecated by now. To circumvent this issue, we retrieved the data from secondary sources as described in Table 2.1.2. We excluded datasets containing less than four sequences. We sampled approximately one third of these test datasets from TreeBASE (Piel et al. 2000) following the dataset curation protocol described in Supplement A. Note that, the BAliBASE sequence sets used for training also, but not exclusively, comprise RV11 and RV12 described in the preceding section.

#### 2.1.3 Entirely Unseen Test Data

We use the PANDIT dataset (version 17.0) (Whelan et al. 2006) to validate the final model on entirely unseen data. PANDIT contains 7738 families of homologous protein domains. For each family, DNA as well as corresponding AA MSAs are available. Note that, for each protein family, PANDIT only provides those protein sequences for which the corresponding nucleotide sequences are *also* available.

We initially removed identical duplicate sequences from the PANDIT MSAs and again, only retain MSAs comprising at least four distinct sequences. Subsequently, we excluded MSAs comprising more than ten masked residues and unaligned the MSAs that passed these two filtering steps. We further, removed unaligned sequence sets comprising sequences shorter than the maximum *k*-mer size of 13 that is being used by our model. Finally, we retained only those sequence sets for which both, the protein *and* DNA sequences were preserved after applying all preceding filtering steps. The resulting dataset comprises 6447 AA sequence sets along with their corresponding DNA sequence sets (Table 3).

**Table 3:**
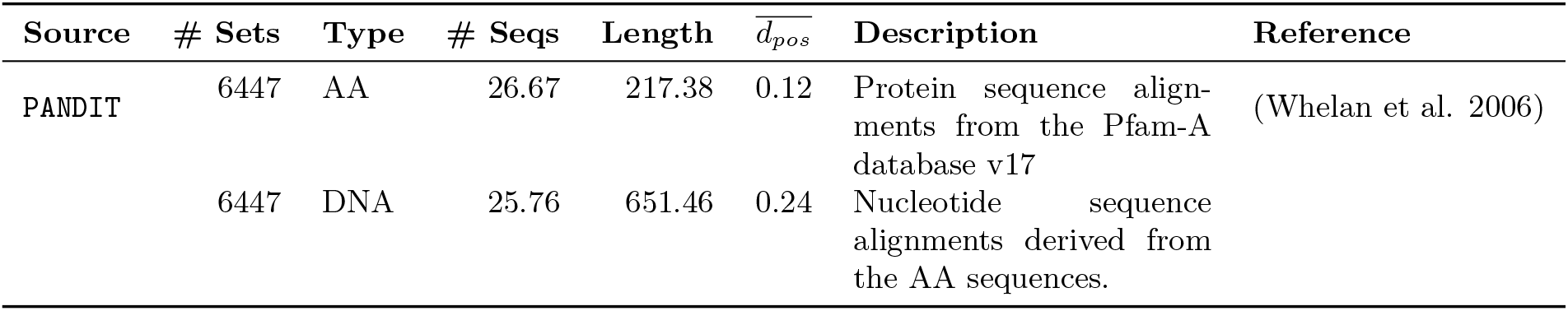
Description of the PANDIT sequence sets used for assessing AlDiScore predictions on entirely unseen data. We use a total of 12 920 sequence sets, divided equally between AA and their corresponding DNA counterparts. Column *#Sets* contains the number of sequence sets per data type source. Column *# Seqs* indicates the mean number of sequences per sequence set. Column *Length* provides the mean length of the longest sequence within each sequence set. The column 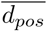 reports the mean value of the reference-free difficulty label 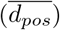 computed on corresponding MSA ensembles (see Section 2.4.1) for each data source.

### 2.2 Generating MSA Ensembles

We already implicitly defined an MSA ensemble as a collection of alternative MSAs inferred on the same single unaligned sequence set via distinct alignment methods and/or alignment method parameter settings. We strive to generate as diverse as computationally feasible (we therefore excluded PRANK (Löytynoja and Goldman 2008), T-Coffee (Notredame et al. 2000) and BAli-Phy (Redelings 2021), for instance) MSA ensembles to maximize the heterogeneity within each ensemble. To this end, we deploy the following widely used and algorithmically diverse alignment methods: Clustal Omega (Sievers et al. 2011), MAFFT (using the FFT-NS-2, G-INS-i, and L-INS-i strategies) (Katoh et al. 2002), MUSCLE v3 (Edgar and Batzoglou 2006), and MUSCLE v5 (Edgar 2022). We consider each of the listed MAFFT strategies and MUSCLE versions as independent MSA approaches and therefore use six distinct MSA algorithms overall. With each of these 6 alignment algorithms we generate 8 MSAs by systematically varying the input parameters according to the settings specified in Table 4. Note that, in Muscle v5 we use the corresponding built-in option for generating an ensemble comprising 8 MSAs. Therefore each ensemble for each unaligned sequence set comprises a total of 48 MSAs (6 algorithms × 8 MSAs per algorithm). Prior to the inference of each of these 48 alignments, we randomly permute the order of the unaligned input sequences to account for the ordering bias that may occur in progressive alignment methods (Landan and Graur 2007). We provide the [ensemblify](https://github.com/MaBody/ensemblify) software package, which enables the systematic and reproducible generation of multiple sequence alignment (MSA) ensembles.

**Table 4:** MSA algorithms and corresponding parameter configurations used for generating MSA ensembles. For each algorithm, we permuted the listed parameters to generate a total of eight MSAs per algorithm. All three disntinct MAFFT algorithms (FFT-NS-2, G-INS-i, L-INS-i) employed the same set of `op` and `ep` parameter permutations.

| Parameter | Value | Reference |
| --- | --- | --- |
| ClustalO v1.2.4 |  |  |
| max-guidetree-iterations | [1, 2, 3, 4] | (Sievers et al. 2011) |
| max-hmm-iterations | [1, 2] |  |
| MAFFT v7.525 (FFT-NS-2, G-INS-i, L-INS-i) |  |  |
| op | [1.1475, 1.4025, 1.6575, 1.9125] | (Katoh et al. 2002) |
| ep | [0.0922, 0.1538] |  |
| Muscle v3.8.31 |  |  |
| weight1 | ["clustalw", "none"] | (Edgar and Batzoglou 2006) |
| weight2 | ["clustalw", "none"] |  |
| objscore | ["spf", "xp"] |  |
| Muscle v5.1 |  |  |
| replicates | 2 | (Edgar 2022) |
| stratified |  |  |

### 2.3 Quantifying MSA Difficulty

In this Section we initially characterize the desired properties of a measure for quantifying MSA difficulty. Then, we describe the two major categories of measures and scoring functions we considered. Finally, for the sake of a more linear text flow, we provide intermediate results to determine the measure that performs best and that will be used throughout the remainder of this work to quantify and predict alignment difficulty.

#### 2.3.1 Determining the Appropriate Scoring Function

To identify an appropriate scoring function for quantifying the MSA difficulty on a given MSA ensemble, we need to determine the function that minimizes the difference between (i) the absolute difficulty as calculated between the MSAs in the ensemble and the reference MSA and (ii) the relative difficulty as calculated from the MSAs in the ensemble only, in absence of a reference MSA. We use the term “reference-based score” synonymously with “absolute difficulty,” and the term “reference-free” synonymously with “relative difficulty.” This is necessary as the latter constitutes the standard use case, that is, reference MSAs are generally not available. To this end, we take as reference our BAliBASE subset (see Section 2.1.1) and use the Pearson correlation coefficient (*r*) and the Earth’s Movers Distance (EMD) to measure how well the relative difficulty corresponds to the absolute difficulty for each scoring function and metric we consider.

Overall, we test two distinct categories of scoring approaches, pairwise and set-based scoring functions (see subsequent Sections for details and Table 5 for an overview). For all pairwise scoring functions, we calculate the absolute difficulty by computing the average over the similarity scores between each MSA in the ensemble and the BALiBASE reference alignment. We compute the corresponding relative difficulty by averaging over the similarity scores of all MSA pairs in the ensemble as outlined in Figure 1A. For the set-based scoring functions, we calculate the absolute difficulty by averaging over the dispersion scores of bi-ensembles that comprise the reference MSA and one of the MSAs from the MSA ensemble at a time over all MSAs in the ensemble. We compute the relative difficulty by directly computing the set-based scoring functions on the MSA ensemble.

**Table 5:** Overview of MSA scoring methods/distance metrics considered. *Dispersion* and *Column Confidence* were developed by Edgar (2022) in Muscle v5. The pHash-based metric is based on (Zauner 2010) and *d*_SSP_, *d*_seq_, and *d*_pos_ metrics were defined by Blackburne and Whelan (2012). All remaining methods are introduced by us.

| Type | Name | Description |
| --- | --- | --- |
| Pairwise | *Dispersion* | Fraction of differing columns. |
|  | pHash-based | Perceptual hashing of alignment matrix. |
| | $d_{\text{SSP}}$ | Homology set metric. Ignores gaps. |
| | $d_{\text{seq}}$ | Homology set metric. Identical coding of gaps in sequence. |
| | $d_{\text{pos}}$ | Homology set metric. Identical coding of consecutive gaps. |
| Set-based | *Column Confidence* | Fraction of reproduced columns across ensemble. |
| | $\text{Conf}_{\text{Set}}$ | Number of unique entries per replication set. |
| | $\text{Conf}_{\text{Entropy}}$ | Shannon entropy per replication set. |
| | $\text{Conf}_{\text{Displace}}$ | Binned standard deviation of indices per replication set. |

**Figure 1:**
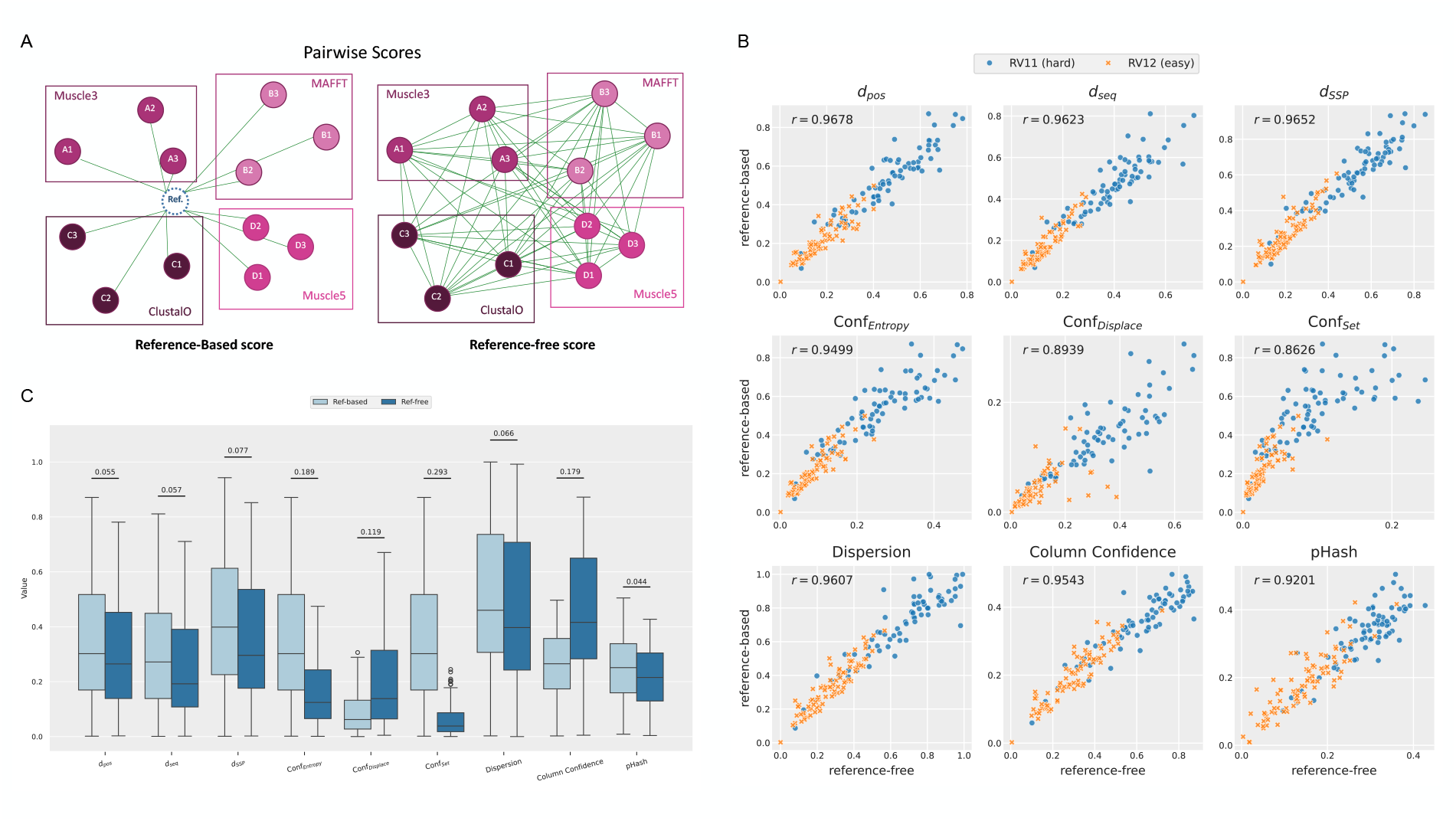
A) Outline of the computation of reference-based, absolute and reference-free, relative pairwise scores. B) Correlation between relative versus absolute difficulties on the BAliBASE RV11 (hard; blue) and RV12 (easy; orange) datasets. All scores are computed on the MSA ensembles generated as described in Section 2.2. Scores in the first row are based on the homology set metrics *d*_pos_, *d*_seq_, and *d*_SSP_ Blackburne and Whelan (2012). The second row contains our newly introduced confusion scores, Conf_Entropy_, Conf_*Displace*_, and Conf_Set_. The third row comprises the *Dispersion* and *Column Confidence* scores from Muscle v5 (Edgar 2022) as well as the pHash-based score. C) Comparison of the absolute (light blue) and relative (dark blue) difficulty distributions computed on the BAliBASE benchmark datasets RV11 and RV12. The number on top of each pair of boxplots indicates the EMD (Earth Movers Distance) between distributions.

#### 2.3.2 Pairwise Scores

In general, we can estimate the inherent dispersion of an MSA ensemble by averaging over pairwise similarity scores or metrics. An overview over the similarity scores we tested is provided in Table 5. The concrete *Dispersion* score we use was introduced by Edgar (2022) but should not be confused with the more general dispersion term we more colloquially use above to derive a notion of difficulty. It relies upon the *Column Score* (Thompson et al. 1999) to quantify the distance between two MSAs. The Column Score captures the proportion of columns that are not replicated in an MSA pair. One may be tempted to use the*Sum-of-Pairs Score* (SPS) Thompson et al. (1999) to quantify MSA similarity or distance. However, as shown by Blackburne and Whelan (2012), SPS is not a true metric because it violates the principles of symmetry and triangle inequality. Thus, apart from the *Dispersion* score, we only consider true metrics as defined by Blackburne and Whelan (2012) to quantify MSA difficulty by computing pairwise MSA distances. However, we decided to omit *d*_evol_ from our analysis, as previous work has demonstrated that it is proportional to *d*_pos_ (Blackburne and Whelan 2012). Relevant implementation details for the metrics are provided in Supplement B.

The central concept of these metrics is the *homology set*. We outline the details by example of *d*_pos_ which, as we show later-on, is the most adequate metric for calculating MSA difficulty.

Let a sequence *S* be an ordered list of characters from a finite alphabet ∑ with length |*S*| = *n* such that *S*[*i*] ∈ ∑. An alignment *A* of sequences *S*_*i*_, *i* ≤ *M* is an extension of these sequences with gap characters such that |*S*_*i*_| = |*S*_*j*_| for all *i, j* ≤ *M*. We denote the aligned sequences as *Ŝ*. A column *A*_*i*_ consists of the characters of *Ŝ*_*j*_ [*i*], the character in position *i* from *Ŝ*_*j*_ ∈*A*.

The distance between MSAs *A* and *B*, denoted by *d*_*pos*_, is defined as

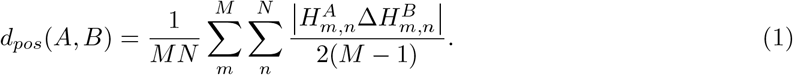

Here, 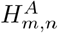 is the positional *homology set* of MSA *A*, sequence index *m*, and character at position *n*, and Δ is the symmetric difference. In other words, if, for two aligned sequences *Ŝ* and 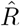 the character at *S*_*i*_ maps to the same position as the character *R*_*i*_, then the *homology sets H*_*S,i*_ and *H*_*R,i*_ are identical. Then, the distance between MSAs *A* and *B* is the average size of the symmetric difference between the homology sets over all site and sequence pairs. See Blackburne and Whelan (2012) for a detailed discussion and formal definition of the relevant *homology set* based metrics we use here: *d*_pos_, *d*_SSP_, and *d*_seq_. Apart from these *homology set*-based pairwise metrics, we also consider the fundamentally distinct pHash-based metric that leverages a Perceptual Image Hash function (Zauner 2010) to create a compact representation of the MSA. The distance between two alignments is computed as the Hamming distance between their perceptual hashes, normalized by the hash length.

As already mentioned, we use these pairwise similarity metrics to compute the average pairwise similarity over all MSA pairs *A*^(*i*)^ and *A*^(*j*)^ for all *i, j* ∈ {1, …, *I*} with *i* ≠ *j*, where *I* is the size of the ensemble. For *d*_pos_ this value is defined as:

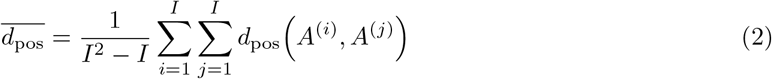

#### 2.3.3 Set-based Scores

Set-based scores can be directly computed over all MSAs by exploiting the set structure of the ensemble. This concept was originally introduced by Edgar (2022), who formally defined the *Column Confidence* score. The column confidence score quantifies how well columns are reproduced across an MSA ensemble. A column that is present in only one MSA of the ensemble obtains a score of 0, while a perfectly replicated column across all MSAs in the ensemble obtains a score of 1 (Edgar 2022). In our setting, we directly subtract the mean *Column Confidence* score from 1.0 such that it corresponds to our difficulty scale where 0 is easy- and 1 is difficult-to-align. Also note that for calculating the relative difficulty we compute these set-based scores directly over the entire MSA ensemble while for obtaining the absolute difficulty we average the score over the bi-ensembles that always contain the reference MSA from BAliBASE (RV11 and RV12) and one of the MSAs from the ensemble.

In addition, we experimented with three distinct variants of a confusion score (Conf_Set_, Conf_Entropy_) which constitutes a novel set-based approach for measuring variation in an MSA ensemble. Since, as already mentioned, the pair-wise *d*_pos_ metric performed best, we describe this score in Supplement C.

#### 2.3.4 Selecting a Metric for Quantifying MSA Difficulty

The intermediate results of our assessment for identifying the most adequate scoring function with respect to its correspondence between the induced absolute (reference-based) and relative (reference-free) difficulty are provided here as they are pivotal for outlining the development of the difficulty prediction approach.

Figure 1B shows the Pearson correlation between the relative and absolute difficulty scores for all metrics listed in Table 5. The absolute and relative difficulties are computed on the MSA ensembles (each comprising 48 MSAs as shown in Table 4) and also comprise the BAliBASE reference MSA for the absolute difficulty calculations as described above. Higher correlation coefficients indicate a better match between the absolute reference-based and relative reference-free difficulty. The highest coefficient of 0.968 is attained by *d*_pos_. Further, all three homology-based set metrics (first row in Figure 1B) score best. In addition, all methods considered yield correlations above 0.85. This indicates that the absolute difficulties can be accurately approximated via relative difficulties in general. Furthermore, all scoring methods considered confirm the prior expectation that BAliBASE RV11 sequence sets are substantially more difficult to align than BAliBASE RV12 sequence sets.

Ideally, the relative difficulties shall not simply correlate with, but directly estimate the values of the absolute difficulties. Figure 1 provides an overview of the respective relative and absolute difficulty value distributions per scoring function we consider.

Overall, the relative difficulties tend to underestimate the absolute difficulties. The homology set metrics *d*_pos_ and *d*_seq_ (Blackburne and Whelan 2012), as well as the Dispersion score (Edgar 2022) and the pHash-based metric, show relatively strong distributional alignment (EMD < 0.07). The *Dispersion* score (Edgar 2022) yields the largest overall difficulty scores. It is the only scoring method that attains a maximum difficulty of 1.0. This is suboptimal for the data at hand, because the *Dispersion* score will exhibit limited discriminative power on very hard datasets. Conversely, the pHash-based metric barely attains scores exceeding 0.5, which indicates that it is not to scale.

While Conf_Entropy_ and *Column Confidence* (Edgar 2022) exhibit high correlations of *r* = 0.95 in Figure 1, their corresponding value distributions in Figure 1 exhibit pronounced scale differences.

Thus, taking into account the correlation coefficient (*r* = 0.9678) in conjunction with the similarity between the absolute and relative difficulty value distributions (EMD= 0.055), the most appropriate relative difficulty metric is *d*_*pos*_. Therefore, we henceforth compute the relative MSA difficulty as the average pairwise distance, denoted as 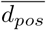, over all MSAs in an ensemble using (2).

### 2.4 Predicting MSA Difficulty

Evidently, computing an MSA ensemble as outlined in Section 2.2 to merely calculate its difficulty is computationally not practical. Ideally, we desire to predict an MSA’s difficulty directly from the unaligned sequence prior to inferring any MSA. Given the difficulty definition derived in the preceding Section, we can now formulate this difficulty prediction task as a supervised regression task. Our objective is to predict MSA difficulty on a continuous scale ranging from 0.0 (easy) to 1.0 (difficult). To this end, we initially compute difficulty labels that reflect the relative MSA difficulty of the training data based on MSA ensembles comprising 48 MSAs per sequence set. Subsequently, we train the `AlDiScore` regression model based upon features that are exclusively computed on the unaligned sequence set.

#### 2.4.1 Label Generation

We compute relative MSA difficulty labels (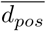 (2)) over all 48 MSAs per MSA ensemble contained in our training data (see Section 2.1.2). We generate MSA ensembles as described in Section 2.2. Note that, even for the BAliBASE datasets (RV11 and RV12) used for assessing the adequacy of distance metrics that are contained in the training data collection, we nonetheless calculate and use the relative difficulties.

#### 2.4.2 Feature Generation

For predicting MSA difficulty, we calculate two main feature categories on the unaligned sequence sets to train the `AlDiScore` regression model. Most features are inspired by those proposed by Trost et al. (2023). However, some phylogenetically inspired features from that work were deliberately excluded here because they do not provide additional predictive power.

We derive a substantial feature subset (PSA-based feature category) from multiple pairwise sequence alignment (PSA) sets generated on the original unaligned sequence set using the Parasail library (Daily 2016). We parametrize the dynamic programming algorithm according to the respective sequence data type. For DNA sequence sets, we apply the DNAfull substitution matrix with a gap opening penalty of 10 and a gap extension penalty of 1. For amino acid (AA) sequence sets, we use the BLOSUM62 substitution matrix with a gap opening penalty of 10 and a gap extension penalty of 1.

For most of our fundamental features, we compute 18 derived features via summary statistics, including the minimum, maximum, mean, standard deviation, inter-quartile range (p80–p20), and 13 percentiles (1, 5, 10, 20, 30, 40, 50, 60, 70, 80, 90, 95, and 99) (Table 6).

**Table 6:** Summary of the features we compute on the unaligned sequence sets. From the majority of the raw feature values, we derive multiple features using 18 summary statistics (minimum, maximum, mean, standard deviation, inter-quartile range (p80–p20), and 13 percentiles).

| Concept | Feature Group | # Features | Description |
| --- | --- | --- | --- |
| Transitive Consistency on PSA triplets | psa_tc | 90 | Transitive consistency of PSA triplets. For each triplet we obtain mean, std, min, max and p50. |
| PSA Length | psa_score_ratio | 18 | PSA score scaled by the minimum sequence length. |
|  | psa_stretch_ratio | 18 | Ratio between max. sequence length and alignment length. |
| PSA Gap | psa_gap_len | 36 | Features (mean, std) based on the lengths of gap regions. |
|  | psa_gap_ratio | 18 | Number of gaps divided by total number of characters. |
| PSA Jensen–Shannon Div. | 4-mer_js | 18 | Pairwise Jensen–Shannon divergence of 4-mer distributions. |
|  | 7-mer_js | 18 | Pairwise Jensen–Shannon divergence of 7-mer distributions. |
|  | char_js | 18 | Pairwise Jensen–Shannon divergence of character distributions. |
|  | hpoly_js_len | 18 | Pairwise Jensen–Shannon divergence of homopolymer length distributions. |
|  | hpoly_js_count | 18 | Pairwise Jensen–Shannon divergence of homopolymer count distributions. |
| Seq. Shannon Entropy | 4-mer_ent | 18 | Shannon entropy of 4-mer distributions. |
|  | 7-mer_ent | 18 | Shannon entropy of 7-mer distributions. |
|  | 10-mer_ent | 18 | Shannon entropy of 10-mer distributions. |
|  | 13-mer_ent | 18 | Shannon entropy of 13-mer distributions. |
|  | char_ent | 18 | Shannon entropy of character distributions. |
| gap % | lbgp | 1 | Lower bound on the gap percentage |

In the following we describe the PSA-based and PSA-agnostic feature categories in more detail.

##### PSA-based Features

We compute the majority of features on PSA sets that are obtained by sampling from the original unaligned sequence set. Note that this sampling-based feature generation introduces some stochasticity in the feature generation process. Although PSA sets can, in principle, be arbitrarily large, we subsample sequence triplets (*n* := 3). Larger tuples (e.g., *n* := 4) increase the number of pairwise values per tuple, which may reduce the per-tuple variance. However, this also limits the total number of sampled tuples for a given CPU time limit for feature computation. In practice, we did not observe any prediction quality improvement when using 4-tuples instead of 3-tuples.

By sampling multiple PSA triplets, we can explicitly control the trade-off between output variance and computational cost for feature computation. By default, `AlDiScore` uses up to 100 triplets (corresponding to at most 300 unique PSAs). This upper limit can be manually specified using the `--max-samples` command line parameter when running `AlDiScore`. For training the model, we employed a higher upper bound of 333 triplets. The higher bound during training means we compute up to 1000 unique PSAs per sequence set, reducing noise due to sampling variance.

We use the PSA triplet structure to compute features for the `psa_tc` feature group based on the notion of Transitive Consistency (Chang et al. 2014). The Transitive Consistency Score is an alignment reliability metric that quantifies the extent to which aligned positions remain consistent across all possible PSA triplets (Chang et al. 2014). A comprehensive description of its implementation in the `AlDiScore` library is provided in Supplement D. For each triplet, we compute the minimum, maximum, mean, median, and standard deviation of this transitive consistency measure. These statistics are subsequently aggregated over all triplets to obtain the previously defined set of 18 summary statistics, resulting in the 90 features associated with the `psa_tc` feature group reported in Table 6.

Additional PSA-based features rely on measuring the length and number of gaps per PSA. The `psa_score_ratio` feature returns the optimal local alignment score using the Smith–Waterman algorithm (Smith and Waterman 1981) normalized by the length of the shorter of the two sequences. The `psa_stretch_ratio` is computed as the ratio between the length of the longest sequence and the full length of each PSA. The `psa_gap_ratio` quantifies the proportion of gaps in a PSA, computed as the number of gaps divided by the total number of characters in the PSA. The `psa_gap_len` measure quantifies the lengths of gap regions. For each PSA we quantify both, the mean, and the standard deviation across all gaps. All gap- and length-related PSA features are subsequently aggregated across all unique PSAs using the aforementioned 18 summary statistics.

Finally, we compute pairwise Jensen–Shannon divergence on all unique PSA. The Pairwise Jensen-Shannon divergence quantifies how different the character usage is between sequences. Higher divergence values indicate greater dissimilarity, which may be associated with harder-to-align sequences. We compute the pairwise Jensen-Shannon divergence of single-character distributions, *k*-mer distributions with *k* ∈ {4, 7}, and homopolymer count as well as length distributions across all PSA alignments. Larger *k* values for the *k*-mers increased the computational cost without improving prediction performance. For each of these divergence measures, we again compute the aforementioned 18 summary statistics to characterize their distributions.

##### PSA-agnostic Features

We also use PSA-agnostic features to characterize other properties of the unaligned sequence set.

We compute these features based upon the Shannon entropy to quantify sequence complexity and variability. Thus, higher entropy values correspond to more diverse residue usage and may be associated with increased MSA difficulty. We compute the Shannon entropy for each individual sequence in the sequence set by using single-character distributions as well as *k*-mer distributions with *k* ∈ {4, 7, 10, 13}. The specific values are a product of iterative trials during development. For each entropy measure, we again calculate the aforementioned 18 summary statistics.

Beyond these entropy-based measures, we also calculate the lower-bound gap percentage (`lbgp`) of the sequence set, that is, the minimal proportion of gap characters required to construct an MSA. This measure is defined as one minus the ratio of the mean sequence length over the maximum sequence length in the sequence set.

#### 2.4.3 Model Training

We trained a LightGBM regression model (Ke et al. 2017) to predict the relative MSA difficulty for a given unaligned sequence set. To this end, we compute the features outlined in the preceding section on each unaligned sequence training set and subsequently employ a repeated as well as nested cross-validation (CV) strategy to determine the optimal model configuration. See Figure 2 for an overview of the training process.

**Figure 2:**
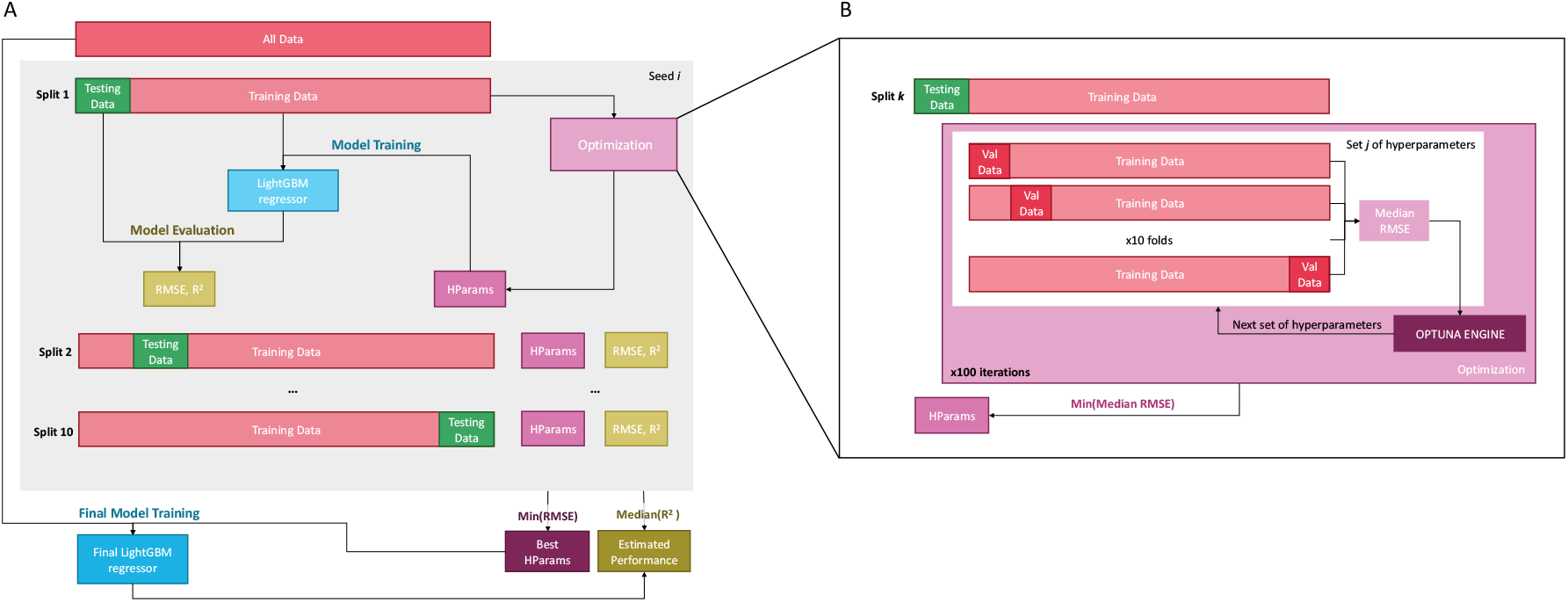
Repeated nested cross-validation pipeline for training and evaluating `AlDiScore`. **(A)** Repeated outer 10-fold cross-validation (CV) across 10 random number seeds (*i* ∈ {1, …, 10}) to accommodate for the stochasticity in the feature generation process, yielding a total of 100 outer splits. For each outer split, we hold out one fold as the testing set (green) while the remaining nine folds constitute the training set (light pink). We optimize Hyperparameters (HParams, purple) in an inner loop. We train the final model on the full dataset using the best hyperparameters across all outer splits (lowest RMSE). We summarize its performance via the median *R*^2^ = 0.945 (10–90% quantiles: 0.939–0.951). **(B)** Inner 10-fold CV for hyperparameter optimization using Optuna (Akiba et al. 2019). We evaluate each hyperparameter configuration across the 10 inner validation folds over 100 hyperparameter search iterations. For each outer fold, we select the hyperparameter configuration that yields the minimum median RMSE across all iterations.

Note that our feature generation process is non-deterministic as it entails random sampling of PSA triplets. Therefore, we partitioned the dataset into the common 10 folds using stratified quantile binning of the continuous target variable to preserve the label distribution across folds, and repeated this procedure 10 times using distinct feature-generation random number seeds to account for the aforementioned inherent stochasticity in the feature generation process. This nested approach therefore yields a total of 100 outer splits (Figure 2A). For each outer split, one fold is held out as test set, while the remaining nine folds constitute the training set.

For each of the 100 outer splits, we perform an inner 10-fold CV to optimize the model hyperparameters via Optuna (Akiba et al. 2019). We evaluate a total of 100 hyperparameter configurations and train each with a fixed number of 1200 estimators and subsequently assess them across all 10 inner validation folds. At the end of the optimization process, we select the configuration that yields the lowest median for the root mean squared error (RMSE) across the inner stratified folds (Figure 2B).

After hyperparameter selection, we retrain the model on the entire outer training set using the selected hyperparameters and conduct an evaluation on the corresponding outer test set. We repeat this procedure for all outer folds and all repetitions to obtain one RMSE value and one *R*^2^ coefficient per outer split.

We train two final `AlDiScore` models (v1.0) on the full AA and DNA data collections, respectively using the hyperparameter configuration that yielded the corresponding lowest RMSE for each data type (Supplementary Table 2).

## 3 Results and Discussion

### 3.1 Experimental Setup

#### 3.1.1 Model Evaluation Experiments

We summarize the performance of our predictive model by aggregating over the RMSE, MAE, correlation, and *R*^2^ coefficients obtained from the 100 outer test folds across all repetitions. We report the corresponding means, along with the 10th and 90th percentiles.

We further also evaluate our final `AlDiScore` model on the yet completely unseen PANDIT dataset (Whelan et al. 2006), which was neither used for model training nor for hyperparameter optimization. Therefore, we initially compute the relative MSA difficulty labels by inferring MSA ensembles as described in Section 2.4.1 and subsequently calculating the corresponding 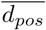 score. Then, we use `AlDiScore` to predict the MSA difficulty on the corresponding unaligned PANDIT sequence sets. We assess the performance of our model on this completely unseen data by computing the correlation between the predicted `AlDiScore` scores and the ground truth relative MSA difficulty scores from the MSA ensembles.

#### 3.1.2 Runtime Analysis Experiments

All runtime experiments for comparing `AlDiScore` runtimes with the runtimes required to calculate the ground truth labels were conducted on a 2-socket machine equipped with two Intel(R) Xeon Gold 6148 CPUs running at 2.40GHz. The system has 40 physical cores (80 threads) and 754GB RAM. To ensure comparability of runtimes, all tools were executed sequentially, that is, on a single physical core.

For these runtime experiments we sampled 35 representative sequence sets based on their size from the PANDIT AA dataset. We define the size as the number of sequences times the maximum sequence length per sequence set. For each sequence set, we measured the times required (i) to generate the MSA ensemble, (ii) to compute the corresponding ensemble-based relative difficulty score 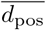 (2), and (iii) to predict the MSA difficulty score on the unaligned sequence set. We add (i) and (ii) to obtain the overall time required for quantifying MSA difficulty via explicit ensemble generation.

We also dissect the prediction time by measuring the contribution of the PSA computations with respect to the overall prediction time. In this experiment, we use the default `--max-sample=100` setting. Hence, we randomly sample at most 100 triplets (less if the sequence set contains but a few sequences) per dataset.

### 3.2 MSA Difficulty Prediction with AlDiScore

As stated above, our training dataset comprises 9651 unaligned sequence sets from a variety of diverse data sources (Table 2.1.2). Note that we distinguish between aligning protein versus DNA sequences. Therefore, we trained a dedicated amino acid (AA) model and a separate, dedicated DNA model. The final `AlDiScore` models predict the MSA difficulty on a scale from 0 (easy) to 1 (difficult).

We initially assess the performance of our model across the 100 outer folds. The median coefficient of determination (*R*^2^) across all folds for the AA model is 0.958 (10th–90th percentile: 0.950 – 0.965). The *R*^2^ for the DNA model is slightly lower, 0.948 (10th–90th percentile: 0.942 – 0.961) as shown in Figure 3B).

**Figure 3:**
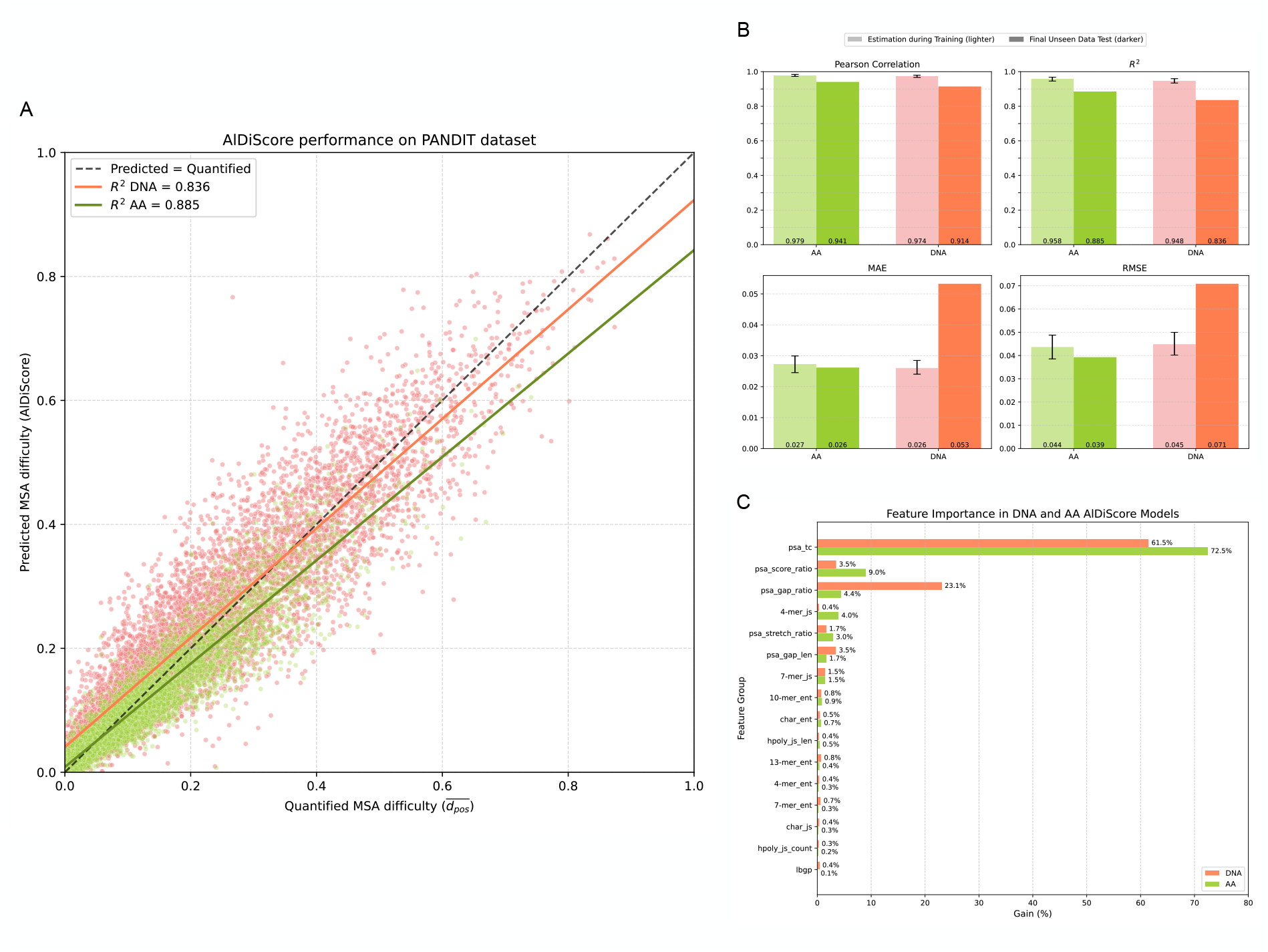
`AlDiScore` models performance and feature importance. Coral refers to DNA and green to amino-acid data. A) Comparison between predicted (`AlDiScore`) and relative 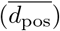 values on the entirely unseen PANDIT test set (Whelan et al. 2006). The *R*^2^ coefficient is 0.84 for DNA data and 0.89 for AA data. B) Performance metrics (RMSE, MAE, correlation, and *R*^2^) for AA and DNA models on training and testing data. The light color columns refer to the summarized performance across 100 outer folds and is reported as the mean with whiskers indicating the 10th–90th percentiles. The darker color columns correspond to our evaluation on the PANDIT dataset. C) Relative feature importance (gain, %) for DNA and AA models across feature groups. Gains are aggregated at the feature-group level. Notice that some groups have more features than others (Table 6).

On the completely unseen PANDIT dataset (Whelan et al. 2006), the model attains an *R*^2^ coefficient of 0.86 for the AA model and of 0.84 for the DNA model (Figure 3A) and a Spearman Rank coefficient of 0.95 and 0.92 for the AA and DNA model respectively. As expected, these values are lower than those obtained from the training process. We nonetheless consider them to be sufficiently accurate to assess the MSA difficulty and inform all subsequent data analysis steps.

#### 3.2.1 AlDiScore Feature Importance

Feature importance differs substantially between the DNA and AA models. While for DNA as well as for AA data, `psa_tc` (transitive consistency of pairwise alignments) exhibits the highest importance, its importance is higher for AA (72.5%) compared to DNA (61.5%) data. This indicates that the consistency-based signal in the sequence set is particularly informative for predicting the difficulty of AA alignments. In contrast, the DNA model relies substantially more on gap-related features, especially on the `psa_gap_ratio` (23.1%), which is only of minor importance in the AA model (4.4%). The prediction for AA data relies more strongly on the `psa_score_ratio` group (9.0%). This suggests that the gap structure plays a more substantial role in DNA alignment difficulty.

Overall, these pronounced differences in feature gains indicate that DNA-based difficulty predictions are dominated by gap patterns and sequence variability at the nucleotide level, whereas AA-based predictions more strongly rely on pairwise alignment consistency. These observations also serve as an *a posteriori* rationale for training distinct AA and DNA models.

### 3.3 Runtime Analysis

We quantified the time required to compute the relative MSA difficulty versus predicting this difficulty via `AlDiScore` across different input size bins. Each bin contains 5 distinct unaligned AA sequence sets from PANDIT.

For all dataset sizes, `AlDiScore` runtimes are substantially faster than the combined runtimes for MSA ensemble generation and computing the relative difficulty (Figure 4A). On small datasets, `AlDiScore` is at least 5 times faster. On large datasets it attains a 700-fold speedup, that is, almost three orders of magnitude. This trend reflects the unfavorable scaling behavior of MSA ensemble generation, whereas the prediction model maintains comparatively low computational complexity.

**Figure 4:**
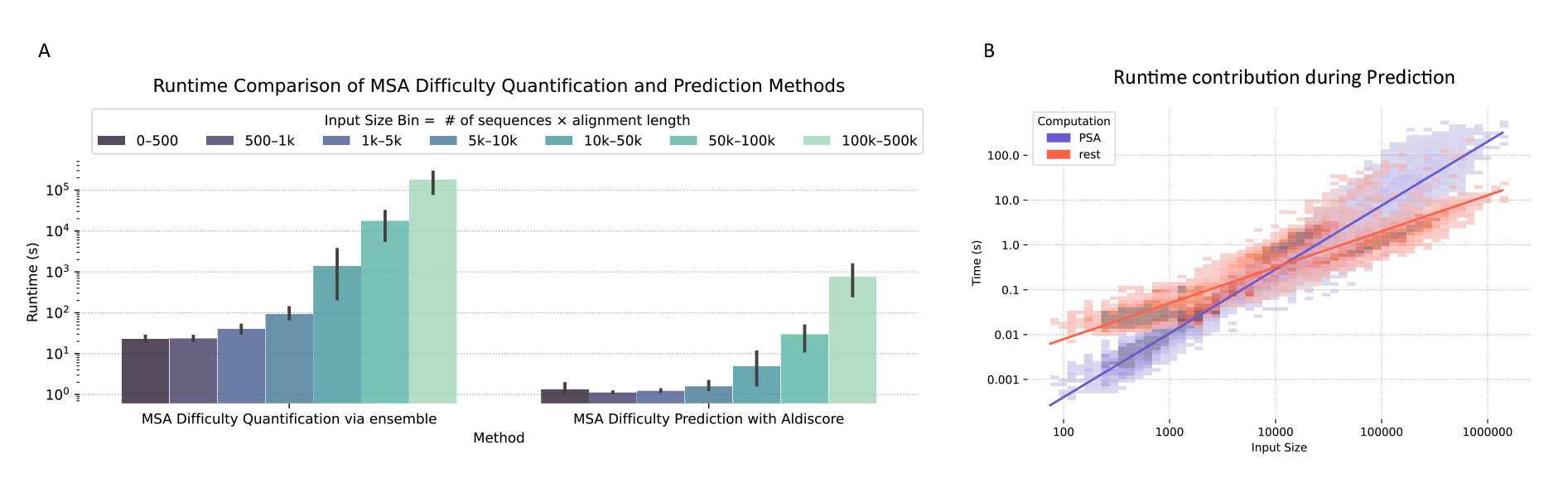
A) Runtime comparison between explicit MSA ensemble generation and label computation via 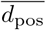 (2) and corresponding `AlDiScore` prediction times. Note that the y-axis is displayed on a logarithmic scale. The x-axis represents input sequence sets grouped into size bins. The bins have been chosen such that their relative proportions approximate a logarithmic scale. The size is defined as the number of sequences times the maximum sequence length. We processed a sample of 35 datasets (5 per bin) using PANDIT AA data on the same hardware and measured sequential execution times on a single physical core. B) Runtime contribution of the PSA computation (blue) compared to the remaining feature extraction steps (red) during the MSA difficulty prediction using `AlDiScore` with default parameters. Both axes are shown on a logarithmic scale. PSA features are computed using the default setting of `--max-sample=100`, that is, at most 100 triplets are randomly sampled per dataset (eventually less on small datasets with fewer sequences).

In Figure 4B we further dissect the prediction time. The results show that the PSA generation step dominates the prediction time, while computing the remaining features is computationally inexpensive. As expected, PSA runtimes scale approximately linearly with the number of sampled triplets (controlled by `--max-sample`). Additionally, there is no correlation between MSA difficulty and input size (Figure 5, *r* = −0.0203). Hence, generating an MSA ensemble is not necessary by default on large datasets as they may be easy-to-align (e.g., the SARS-CoV-2 datasets from Morel et al. (2021) obtained a difficulty of 0.0).

**Figure 5:**
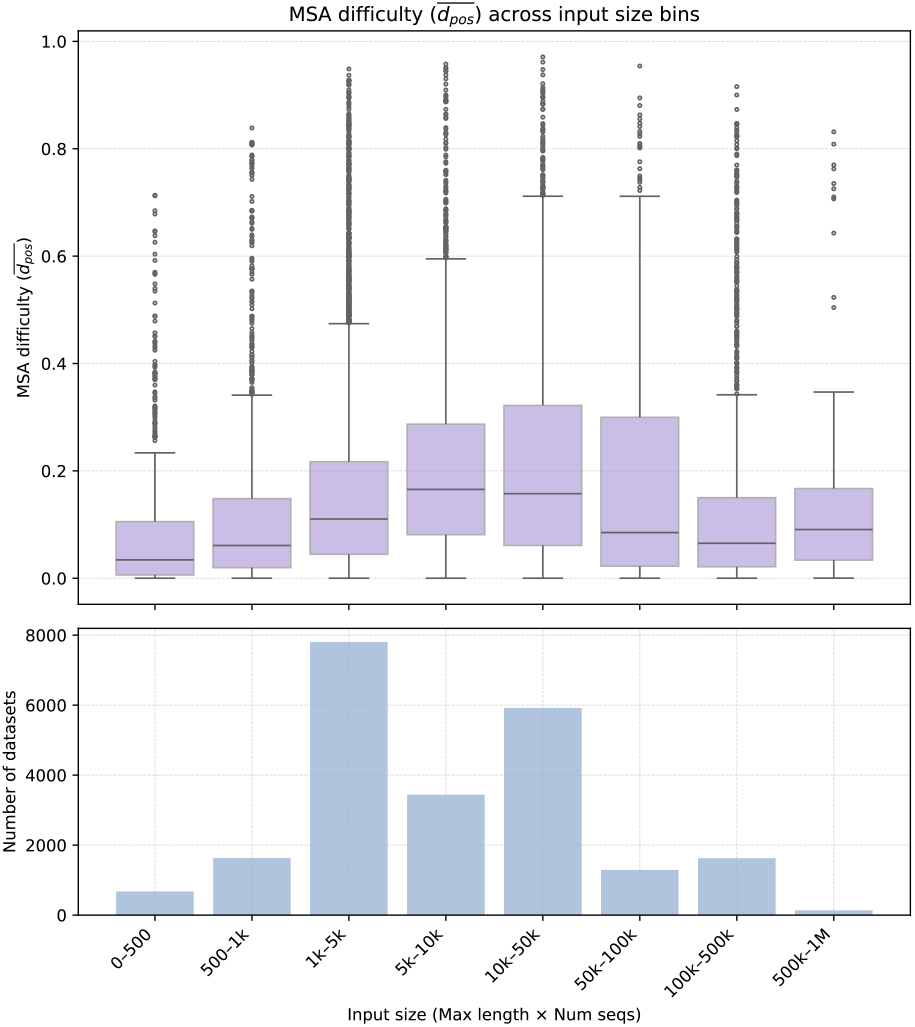
Comparison between datasets size and MSA difficulty. The correlation coefficient between input size (Max length x Num seqs) and MSA difficulty (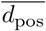 is −0.0203. Upper box-plots represent the MSA difficulty distribution across different dataset sizes. The bottom plot represents the number of datasets in each input size bin.

### 3.4 MSA difficulty of DNA versus corresponding AA datasets

We also studied the relationship between the MSA difficulty labels computed on the explicit MSA ensembles via 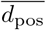 (2) for AA sequence sets and their corresponding DNA sequence sets (i.e., MSA difficulty for the same taxa) using the PANDIT dataset (Figure 3.4). We observe a strong positive correlation with a determination coefficient of *R*^2^ = 0.831. This indicates that sequence sets that are difficult-to-align at the DNA level, also tend to be difficult-to-align at the AA level. The fitted linear model (*y* = 0.58*x* + 0.01) exhibits a slope that is substantially below 1.0. Hence, AA sequence sets are consistently easier-to-align than their DNA counterparts. The higher difficulty of DNA alignments is plausibly explained by the increased variability at the nucleotide level which introduces additional ambiguity during alignment, making DNA sequences 1.7 times harder to align than their corresponding amino acid sequences. In this context, Dessimoz and Gil (2010) observed fewer differences among programs when aligning AA, than when aligning nucleotides sequences and concluded that current alignment packages align amino acids more accurately than nucleotides.

### 3.5 Relationship of Alignment Difficulty with Phylogenetic Difficulty

We show the relationship between the MSA difficulty and the corresponding average phylogenetic difficulty over all MSA per ensemble in Figure 7. Overall, we observe no clear correlation between the two difficulty measures (*r* = −0.07). Hence, the MSA and the phylogenetic difficulty are predominantly independent of each other. While a slight negative trend is present, its magnitude is very small and does therefore not support a strong inverse relationship.

Intuitively, one might expect that sequence sets that are difficult-to-align will also be difficult to phylogenetically resolve. However, the optimization criteria deployed in alignment methods versus phylogenetic inference methods differ. MSA methods strive to minimize differences between sequences by identifying homologous positions, whereas phylogenetic inference relies on exactly those differences to distinguish between alternative tree topologies. Highly divergent sequences can therefore increase alignment uncertainty, while simultaneously providing a stronger phylogenetic signal for the tree inference process. Conversely, highly similar sequences are typically easy-to-align but may yield limited phylogenetic signal, making it harder to confidently resolve evolutionary relationships. A representative example for the latter constellation are SARS-CoV-2 sequences which are generally considered to be easy-to-align (`AlDiScore` = 0.0), but hard to phylogenetically infer (Pythia difficulty = 0.76) (Morel et al. 2021).

Despite this conceptual contrast, the weak correlation we observe here suggests that these effects do not always apply to the datasets we analyzed. In particular, the near-zero correlation indicates that one cannot conclude that difficult-to-align sequences sets are systematically easy (or difficult) for phylogenetic inference. This reinforces the view that MSA difficulty and phylogenetic difficulty represent distinct sources of variation, as uncertainty in phylogenetic reconstruction can arise from factors beyond sequence alignment alone (Dessimoz and Gil 2010). Consistent with this interpretation, Dessimoz and Gil (2010) also reported only a weak association between alignment variability and tree accuracy (*r*_*s*_ = 0.16). Figure 7 nonetheless shows that datasets more commonly exhibit low difficulty for both, the MSA, and the phylogenetic inference task, while cases of simultaneous high alignment and tree inference difficulty are comparatively rare. This is encouraging because it means that inherent inference uncertainty and respective ensembles of MSAs or trees will rarely need to be considered for both, the MSA, and phylogenetic inference steps during a single data analysis.

### 3.6 Determining the optimal Benchmark dataset

Figure 8 shows the MSA and phylogenetic difficulty distributions of all databases used in this study. Alignment benchmark databases are commonly used to evaluate the performance of alignment programs. However, the figure indicates that most benchmark datasets are heavily skewed toward easier alignment difficulties. Because reported alignment accuracy strongly depends on the inherent difficulty of the sequence sets being evaluated, benchmark composition can substantially influence the conclusions drawn about program performance. Failure to account for this effect may lead to misleading interpretations. An example of this issue was observed for DiAlign: although both simulated and structure-based reference alignments suggested that the method had substantially improved across three releases (Subramanian et al. 2008), later analyses by Dessimoz and Gil (2010) using different datasets and tests did not support this conclusion. Shrestha and Adhikari (2022) also observed a 10% increase on their Deep Learning model when including an additional metagenomic database into their training set. These cases illustrate how dataset composition can alter reported performance and emphasizes the importance of considering the difficulty distribution of benchmark datasets when assessing alignment methods.

The overall trend is that for easy alignments, distinct alignment approaches tend to yield highly similar results, whereas in difficult cases, the resulting alignments may vary substantially. Consequently, high alignment accuracy scores may merely reflect that the benchmark is easy, rather than that the alignment method works particularly well.

This issue becomes evident on challenging datasets. The “twilight” subset of SABmark represents one of the most difficult-to-align test sets (Figure 8), and most alignment programs tend to generate low-scoring or incorrect alignments. More generally, difficult datasets frequently expose the limitations of current methods. Aligning non-homologous sequences is often among the most challenging scenarios because proteins that are related through processes beyond simple insertions, deletions, and substitutions do not fit the underlying assumptions underlying assumptions of most alignment algorithms well (Lassmann 2005). Under these conditions, reporting that method A performs marginally better than method B is of limited practical significance, since all methods may fail to recover biologically meaningful alignments (Lassmann 2005).

## 4 Conclusion

In this study, we initially quantify the MSA difficulty as an intrinsic property of an unaligned sequence set. To this end, we deploy scores derived from diversely inferred MSA ensembles deploying widely used but also conceptually distinct MSA tools to capture the variance among MSAs generated from the same underlying unaligned sequence set. The observed ensemble variance serves as an indirect measure of alignment difficulty. In other words, a small variance within the ensemble indicates that the sequences are easy-to-align whereas a high variance indicates that they are difficult-to-align. MSA distance scores between inferred MSAs and corresponding reference MSAs, have been shown to generally serve as strong predictors of alignment error. However, these approaches rely on the availability of high-quality reference MSAs, which are often not available in practice.

Thus, our first contribution was to identify adequate, reference-free measures of alignment difficulty on an ensemble of MSAs. To this end, we evaluated several reference-free scoring methods by assessing how well they recover the difficulty captured by their reference-based counterparts. We conclude that the best approach for quantifying difficulty is 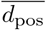 (2). Thus, we quantify MSA difficulty as the average pairwise distance between all MSAs in an ensemble (Herman et al. 2015; Edgar 2022) to capture the relative variance among alternative alignments in an ensemble. As we show, averaging these distances over an ensemble (without a reference MSA) yields a robust estimate of MSA variance, where higher values reflect greater disagreement among MSAs and, consequently, higher alignment difficulty.

Generating such MSA ensembles is computationally expensive. Hence, directly computing difficulty scores for large unaligned sequence sets is computationally not feasible. As we observe no relationship between sequence set size and alignment difficulty, this limitation is particularly relevant for large sequence sets that are easy-to-align, because computing an MSA ensemble may simply not be necessary. This motivates the need for approaches that can predict the difficulty on a given sequence set *a priori*. The development of such a method constitutes the second contribution of this work. As we show, our easy-to-use open source tool `AlDiScore` can accurately predict the alignment difficulty directly on unaligned sequence sets. It also exhibits good accuracy on completely unseen data (*R*^2^ = 0.885 on AA and *R*^2^ = 0.836 in DNA sequences).

Our feature importance analysis advances our understanding of the factors that affect alignment difficulty. Overall, features derived from sampling pairwise alignments constitute the best predictors of MSA difficulty. The difficulty is dominated by the pairwise alignment consistency signal on AA data, while for DNA data, the difficulty is more strongly influenced by gap-related features. We attribute this observation to a higher degree of evolutionary conservation at the protein level as opposed to increased nucleotide sequence variability. We also observe this when directly comparing the MSA difficulties of PANDIT AA sequence sets with those of the corresponding DNA sequence sets which are approximately 1.7 times more difficult-to-align on average.

We further find that MSA difficulty and phylogenetic difficulty show no clear correlation (Pearson correlation coefficient = −0.07). While one might expect highly divergent datasets to be both, hard to align, and hard to phylogenetically infer, our results indicate that MSA and phylogenetic difficulty are determined by distinct characteristics of the underlying data. In general, most MSA tools, strive to *minimize* differences between sequences. Sequence sets that comprise highly divergent sequences therefore tend to be more difficult-to-align. Phylogenetic inference, on the other hand, relies upon exactly those differences to clearly distinguish between alternative tree topologies. That is, the degree of heterogeneity of the sites in an MSA determines the magnitude of differences among likelihood scores for distinct putative candidate phylogenies. If these differences are large, phylogenetic inference tools are better able to identify a “best-known” tree, that is, a clear local maximum. Conversely, if the MSA contains sequences which are highly similar, the likelihood difference between any two putative candidate phylogenies can be (relatively) small such that they become statistically indistinguishable (see (Morel et al. 2021) for a representative example of such a dataset). Therefore, MSA inference tools, in a sense, reduce the information or signal strength that phylogenetic inference methods requires to identify a pronounced peak on the phylogenetic likelihood surface. As a result, sequence sets may be difficult to align but easy to phylogenetically infer or vice versa. This emphasizes the importance of treating MSA difficulty as an autonomous source of variation.

Overall, `AlDiScore` implements an efficient as well as accurate approach for guiding MSA inference. Instead of either relying on a single MSA or incurring the high computational cost of MSA ensemble generation by default, practitioners can now use `AlDiScore` difficulties to inform their analytical setup *a priori*. On an easy sequence set, computing a single alignment via the fastest available MSA method will suffice, whereas a difficult sequence set will require conducting ensemble-based analyses as well as propagating the corresponding uncertainty (ensemble of MSA) to the corresponding downstream analysis step, that is, phylogenetic inference, for instance.

## 5 Future Work

Our new approach to accurately predict MSA difficulty from unaligned sequence sets opens several directions for future research. One may envision developing a fully adaptive phylogenetic inference pipeline that explicitly accounts for the corresponding difficulty at each pipeline stage (e.g., MSA and phylogenetic inference). Overall, such a pipeline will provide an end-to-end framework that dynamically and automatically adapts to dataset-specific characteristics while explicitly accounting for uncertainty throughout the sequence analysis process. Starting on an unaligned sequence set, such a pipeline could use the predicted MSA difficulty to determine whether a fast-to-compute, single MSA is sufficient or whether MSA uncertainty needs to be taken into account. In cases of high MSA difficulty, failing to account for alignment uncertainty can lead to unreliable or misleading results (Wong et al. 2008; Wu et al. 2012). In particularly challenging cases, the difficulty may be so high that no single unambiguous alignment solution exists. A common strategy to address this issue is to generate an MSA ensemble. Such ensembles can then be used for several downstream applications, including performing multiple phylogenetic inferences to assess tree robustness (Edgar 2022), constructing consensus alignments (Wallace et al. 2006), or ranking candidate alignments to identify the most accurate solutions within the ensemble (Lassmann 2005; Shrestha and Adhikari 2022; Serok et al. 2026).

Subsequently, Pythia (Haag et al. 2022) can be used to guide the phylogenetic inference process. In fact, Pythia has already directly been integrated into RAxML-NG v2.0 to determine an appropriate setup of the tree search heuristics ((Togkousidis et al. 2023)). Analogously, we envision that `AlDiScore` could also be directly integrated into the widely-used MSA tools we used here for generating ensembles to fully automate this process and adapt the alignment heuristics being used to the data at hand. Extending the pipeline, difficulty prediction could also be deployed further up- and downstream by taking into account orthology assignment or molecular species delimitation or dating variance, for instance.

Another potential application of `AlDiScore` is in dataset curation and assembly. By applying `AlDiScore` in a leave-one-out fashion, one can quantify the contribution of individual sequences to the overall MSA difficulty. This may allow to identify sequences that disproportionately increase the difficulty and assess whether removing them induces a more stable and reliable high quality MSA. This could contribute to identifying homologous sequences and generating more accurate trees. However, it is important to distinguish between non-homologous genes and those who, while being homologous, have diverged in a non-linear way. The latter genes are of substantial interests and can be used to further improve alignment programs (Wong et al. 2008). This is because current alignment programs may fail to accurately capture the underlying biological relationships between sequences under difficult alignment scenarios (Lassmann 2005). Being able to diagnose challenging alignments (i.e., difficult alignments) constitutes the first step toward improving quality, and “a function used to assess the quality of completed alignments can potentially be used in reverse as an objective function in an alignment algorith” (Lassmann 2005). As noted by Notredame et al. (2000), assessing the quality one a case-by-case basis may be more informative than solely striving to improve average accuracy values. By characterizing the alignment difficulty distribution across benchmarks comprising unaligned sequence sets, one can identify potential biases toward overly easy or difficult sequence sets in the experimental setup. Ensuring that benchmark sequence sets adequately reflect the full real-world difficulty spectrum is essential for conducting fair and meaningful evaluations of alignment tools. As an example, in Figure 8, we depict the MSA and phylogenetic difficulty distribution for the sequence sets used in this study. Therefore, in analogy to Pythia which has already been used for exactly this purpose (see Fig. 6 in (Wiegert et al. 2024)), `AlDiScore` offers the opportunity to systematically assess and improve benchmark datasets used for evaluating MSA methods. Finally, plotting MSA inference accuracy/quality over the difficulty spectrum (as also already conducted for phylogenetic difficulty, see Fig. 2 in (Höhler et al. 2025)) will allow to identify which method, heuristic, or criterion performs best on easy, medium, or difficult sequence sets.

**Figure 6:**
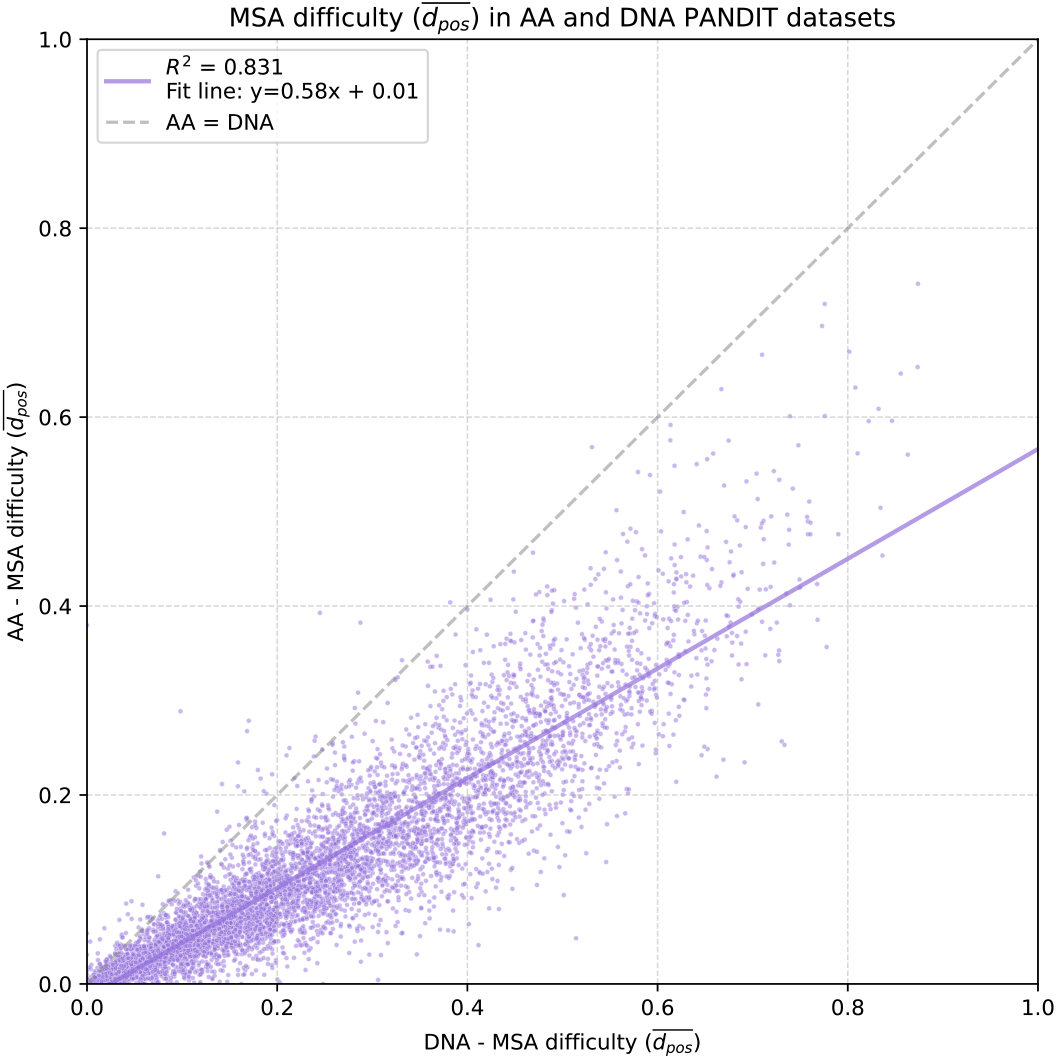
Relationship between the difficulty of aligning DNA and corresponding AA sequence sets from PANDIT (Whelan et al. 2006). The fitted regression line (*y* = 0.58*x* + 0.01) with *R*^2^ = 0.831 indicates that AA sequence sets are generally easier-to-align than DNA sequence sets.

**Figure 7:**
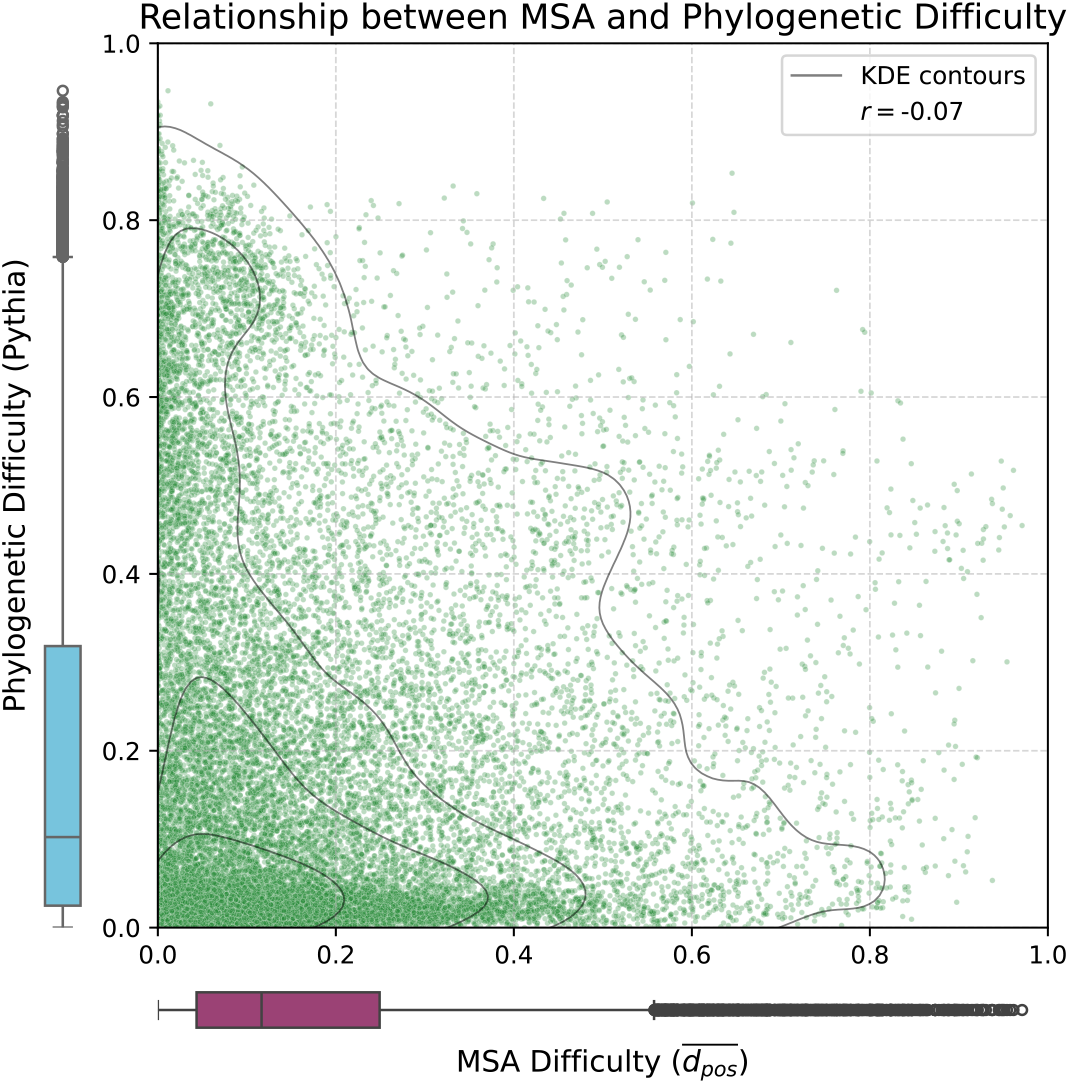
Joint distribution of relative MSA difficulties across all 22 545 sequence sets and corresponding phylogenetic difficulty. We quantify the MSA difficulty using the 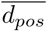 (2) in an explicit MSA ensemble and we compute the phylogenetic difficulty by averaging the difficulties predicted by Pythia v2.0 (Haag et al. 2022) over all MSAs in the respective ensemble. We observe no correlation between phylogenetic and MSA difficulty (*r* = −0.07). The Kernel Density Estimation (KDE) contours indicate regions of higher and lower point density, while the marginal boxplots show the distributions of MSA and phylogenetic difficulties, respectively.

**Figure 8:**
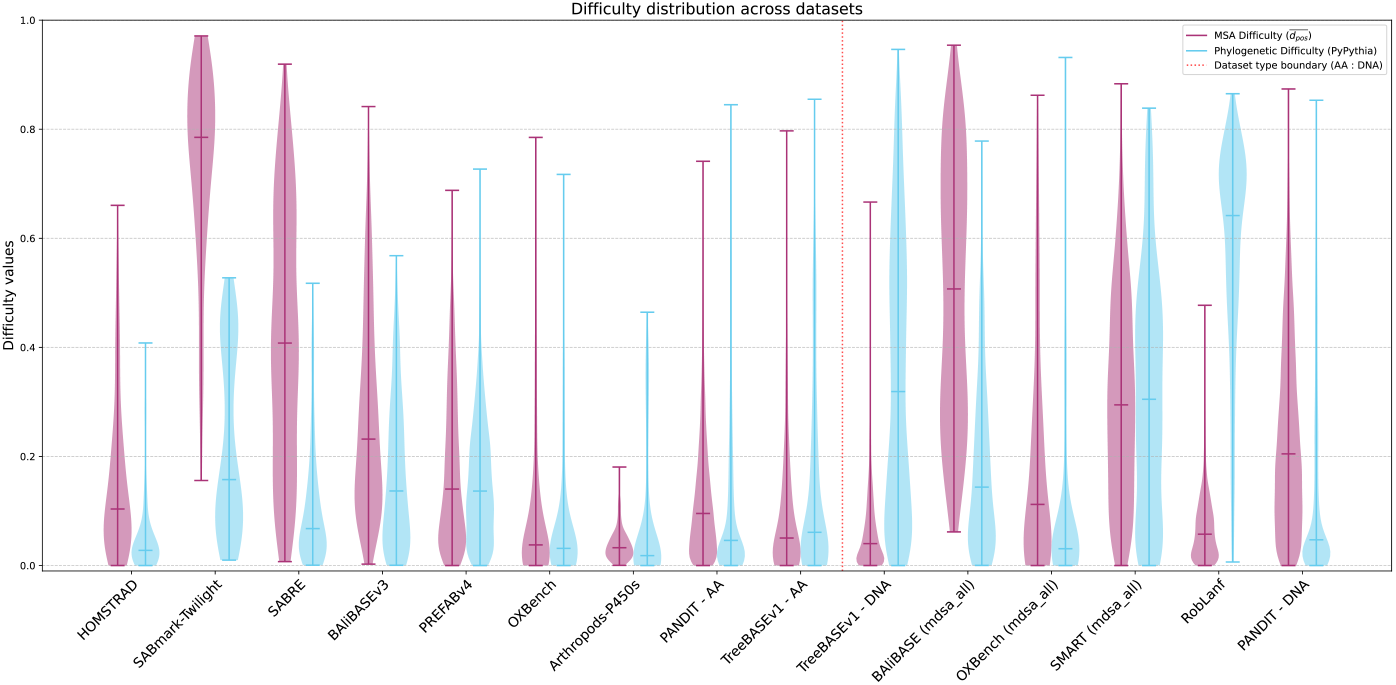
MSA difficulty (dark magenta) and Phylogenetic difficulty (light blue) distribution across databases represented using violin plots. The horizontal lines represent the median, minimum and maximum values on each database. The vertical dotted red line separates amino acid from nucleotide databases. MSA difficulty is quantified using the 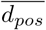 (2). We compute the phylogenetic difficulty by averaging the difficulties predicted by Pythia v2.0 (Haag et al. 2022) over all MSAs in the respective ensemble.

## 6 Code Availability

The implementation and full source code are publicly accessible under GNU General Public License version 3 (GPL v3) at [ensemblify](https://github.com/MaBody/ensemblify) and [AlDiScore](https://github.com/MaBody/aldiscore). The data used in this study is publicly accessible at AlDiScore_data under GPLv3.

