## Supplemental Material for "Quantifying and Predicting the Difficulty of Multiple Sequence Alignment with AlDiScore"

### Supplementary Material

### A TreeBASE curation protocol

We curated TreeBASE MSAs using the steps listed in Table 1. The cleaning step consisted of removing invalid data, such as datasets with empty sequences, invalid characters, or malformed MSA files. Furthermore, to exclude outliers, we removed datasets which:

- Contained a sequence with a length greater than the 95%-ile over all sequences;
- Contained a number of sequences greater than the 95%-ile over all MSAs;
- Or had an Inflation ratio ($I_R$) (see Eq. 1) less than the 5%-ile.

$$I_R(A) := \frac{\sum_{k=1}^{K} |S_k|}{N \times K} \tag{1}$$

where $N$ is the length of the aligned sequences in $A$, and $K$ is the number of sequences in $A$. The inflation ratio measures alignment density as the proportion of residues and the total number of sites in an MSA. After the Outlier Removal step, we constrained the maximum length of the longest unaligned sequence to 10.000 sites to reduce computational costs. The data cleaning and filtering step yielded a total of 7995 sequences sets, 92% of which were DNA. To ensure that large samples are adequately represented in the sample, we applied a weighting scheme proportional to dataset size

| Step | | Remaining | | Total |
|---|---|---|---|---|
| | | AA | DNA | |
| 0 | Start | | | 10404 |
| 1 | Cleaning | 810 | 8486 | 9296 |
| 2 | Outlier Removal | 699 | 7355 | 8054 |
| 3 | Constraints | 640 | 7355 | 7995 |
| 4 | Sampling | 622 | 2497 | 3119 |

Supplementary Table 1: TreeBASE cleaning steps and results

### B Implementation of pairwise scores

The computation of the average pairwise distance between all alignments in an MSA ensemble requires the computation of a distance matrix on each MSA ensemble. With a total of $I$ alignments per ensemble, each alignment has to be compare $I - 1$ times. This problem structure applies to all pairwise scoring methods. By caching intermediate computational states for each alignment, we can exploit the structure to improve performance. Caching allows us to execute a substantial share of the calculations only $I$ times for $I(-1)/2 \in \Theta(I^2)$ comparisons.

To applicate the caching techniques we fully re-implemented $d_{\mathrm{SSP}}$ $d_{\mathrm{seq}}$ and $d_{\mathrm{pos}}$ metrics in Python. For the positional encoding of the *homology sets*, we use an integer system. For each code, we concatenate the indices of sequence and (unaligned) position. To prevent overflow of the positional index, we add zero padding in the middle where necessary. Furthermore, positive integers encode residues, while negative integers encode gaps. Consecutive gaps are encoded with the last residue code before the gap sequence. In Figure B, we give an example for our encoding scheme. The homology set for residue $\hat{S}_2[4]$ corresponds to the character C and the integer code 22. The corresponding homology set is $H_{2,2} = \{-13, 33\}$. For additional examples, we refer to (Blackburne and Whelan 2012).

### C Confusion-based methods

Confusion-based methods compute a score over the entire ensemble by leveraging its set structure. In our implementation, we first calculate a difficulty score for each residue individually and then aggregate

| | $n$ | 1 | 2 | 3 | 4 | 5 | 6 |
|---|---|---|---|---|---|---|---|
| $\sum_{\text{DNA}}$ | $\hat{S}_1$ | A | C | A | - | - | T |
| | $\hat{S}_2$ | - | G | - | C | C | T |
| | $\hat{S}_3$ | A | C | - | C | G | - |
| $d_{pos}$ | $\hat{S}_1$ | 11 | 12 | 13 | -13 | -13 | 14 |
| | $\hat{S}_2$ | -20 | 21 | -21 | 22 | 23 | 24 |
| | $\hat{S}_3$ | 31 | 32 | -32 | 33 | 34 | -34 |

Supplementary Figure 1: Example for the positional encoding to compute $d_{\text{pos}}$.

these values to produce a single score for the entire dataset. We define three variants of the confusion score: $\text{Conf}_{\text{Set}}$, $\text{Conf}_{\text{Entropy}}$, $\text{Conf}_{\text{Displace}}$ (Figure 2). Each variant produces a normalized confusion score per residue, which is averaged across all residues to obtain the ensemble-level score. To do so, we define the *replication set* to represent how a fixed residue is aligned across a MSA ensemble. *Replication sets* are based on the concept of positional homology sets proposed by Blackburne and Whelan (2012). The replication set $R_{l,k,n}$ collects, across an ensemble of $I$ alignments, the residues from sequence $l$ that are aligned to residue $s_{k,n}$ in another sequence $k$, where $n$ denotes the unaligned position in sequence $k$. Replication sets therefore provide a compact representation of the full ensemble information by encoding, for each residue, all alignment-based hypotheses of its homology. The replication set $R_{l,k,n}$ therefore contains, for each alignment in the ensemble, the residue from sequence $l$ that is aligned to residue $s_{k,n}$ of another sequence $k$.

We compute $I$ distinct homology sets for each residue, where $I$ denotes the number of alignments in a given MSA ensemble.

For $\text{Conf}_{\text{Set}}$, $\text{Conf}_{\text{Entropy}}$ we applied the gap-sensitive encoding from $d_{\text{pos}}$. However, for $\text{Conf}_{\text{Displace}}$ we applied the positional encoding from $d_{\text{SSP}}$ to exclude gap positions from all direct comparisons.

These *replication sets* provide a compact representation of the full ensemble information. For each residue, replication sets collect the aligned entries from all alignments, effectively representing all the ensemble-based hypotheses of its homology.

Similarly to homology sets, Gap-sensitive encoding is applied by default to better incorporate gap information.

and capture alignment agreement within individual sequences across the ensemble. The resulting scalar serves as a proxy for alignment difficulty: values range from 0 (easy) to 1 (difficult),

The Confusion Based on Set Cardinality score, $\text{Conf}_{\text{Set}}$, quantifies dispersion by counting how many distinct symbols occur in a replication set. For each residue, the number of unique characters aligned to it across the ensemble is computed and min–max normalized. This measure captures whether aligners disagree, but not how strongly or how unevenly that disagreement is distributed.

Confusion Based on Entropy score, $\text{Conf}_{\text{Entropy}}$, extends $\text{Conf}_{\text{Set}}$, by incorporating the full distribution of distinct characters in each replication set. Instead of relying solely on the number of unique entries, it computes the Shannon entropy of their empirical distribution. The entropy is normalized by its theoretical maximum for a sample of the same size, producing values between 0 and 1.

$\text{Conf}_{\text{Entropy}}$ therefore distinguishes between dominant and evenly split alignment alternatives, reflecting a finer-grained notion of uncertainty.

To capture alignment dispersion beyond residue identity, we introduce the confusion Based on Positional Displacement score, $\text{Conf}_{\text{Displace}}$, inspired by the $\text{SP}_{\text{dist}}$ score of Bawono et al. (2015). $\text{SP}_{\text{dist}}$, measures how much a query alignment actually differs from the reference by taking distances between mismatched residue pairs into account in MSA comparison. Previously defined replication set-based measures, $\text{Conf}_{\text{Entropy}}$ and $\text{Conf}_{\text{Set}}$, only consider whether residues are aligned consistently across an ensemble, ignoring the magnitude of positional deviations. $\text{Conf}_{\text{Displace}}$ addresses this by quantifying the dispersion of column indices within each replication set $R_{l,k,n}$.

Analogous to $\text{SP}_{\text{dist}}$, which evaluates positional distances between aligned residue pairs across alignments with thresholding, $\text{Conf}_{\text{Displace}}$ first computes the standard deviation of the column indices in each replication set, capturing how far residues from different alignments deviate from their mean position. To ensure comparability across residues and prevent extreme deviations from dominating the score, the standard deviations are transformed using a threshold-based binning scheme, following the approach of Bawono et al. (2015). Specifically, through empirical evaluation we defined an exponentially scaled threshold set, $T = \{0, 1, 2, 4, 8, 16, 32\}$, in order to smooth over larger distances. The final $\mathrm{Conf}_{\mathrm{Displace}}$ value for residue $s_{k,n}$ is obtained by averaging the binned displacement scores over all $K - 1$ replication sets corresponding to sequences $l \neq k$. The final $\mathrm{Conf}_{\mathrm{Displace}}$ scores reflects how consistently aligners place homologous residues at similar positions, capturing MSA dispersion due to spatial displacement along the sequence.

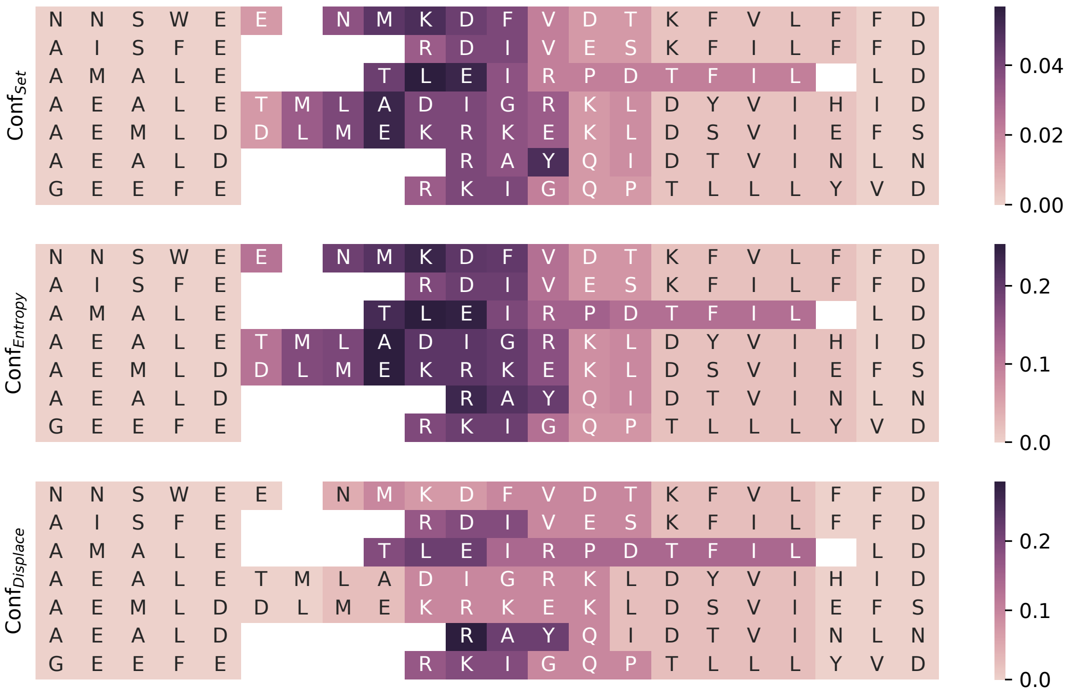


Supplementary Figure 2: Visualization of an example for the confusion score variants $\mathrm{Conf}_{\mathrm{Set}}$, $\mathrm{Conf}_{\mathrm{Entropy}}$, and $\mathrm{Conf}_{\mathrm{Displace}}$. The displayed region corresponds to indices 109–131 of the dataset BB12036 from BAliBASE RV12 (Thompson et al. 2005). Gaps are shown in white. For the residues, darker colors indicate higher difficulty. Theoretically, all scores are bounded in the [0,1] range. The variants are not scaled identically due to different normalization procedures.

### D Transitive Consistency

Our most important feature group **psa_tc** is based on the notion of transitive consistency by Chang et al. (2014). Transitive consistency is a property of a residue with respect to a PSA triplet. The original implementation of transitive consistency (Chang et al. 2014) was not flexible enough for our purposes. Hence, we implemented our own flavor of transitive consistency in the AlDiScore library.

To compute features based on transitive consistency, we first draw three sequences $S_1$, $S_2$, and $S_3$ uniformly at random and compute all three possible PSAs on this sequence triplet. Let $A_{12}$ be the PSA of $S_1$ and $S_2$, obtained via dynamic programming. Let $A_{13}$ and $A_{23}$ be the PSAs for the remaining sequence pairs. Assume that some residue $s_1$ of the first sequence is aligned with residues $s_2$ in $A_{12}$ and $s_3$ in $A_{13}$. Now, we compare whether $s_2$ is also aligned with $s_3$ in $A_{23}$. If so, the PSA triplet is consistent for residue $s_1$. We repeat this comparison for all residues of a PSA and compute the average consistency. For our feature space, we then compute summary statistics on the resulting values.

### E Final Hyperparameter selection

Supplementary Table 2: Hyperparameter settings for the final LightGBM regressors (AA and DNA AIDiScore models).

| | n_estimators | colsample_bytree | colsample_bynode | learning_rate | min_data | num_leaves | lambda_l1 | lambda_l2 | subsample |
|---|---|---|---|---|---|---|---|---|---|
| AA | 1200 | 0.088135 | 0.217743 | 0.037031 | 22 | 43 | 0.000043 | 0.000207 | 0.402253 |
| DNA | 1200 | 0.276943 | 0.467235 | 0.023612 | 11 | 34 | 0.000339 | 0.000328 | 0.888803 |
